# Time-course transcriptome data of silk glands in day 0–7 last-instar larvae of *Bombyx mori* (*w1 pnd* strain)

**DOI:** 10.1101/2024.03.02.582034

**Authors:** Yudai Masuoka, Akiya Jouraku, Takuya Tsubota, Hiromasa Ono, Hirokazu Chiba, Hideki Sezutsu, Hidemasa Bono, Kakeru Yokoi

## Abstract

Time-course transcriptome expression data were constructed for four parts of the silk gland (anterior, middle, and posterior parts of the middle silk gland, along with the posterior silk gland) in the domestic silkworm, *Bombyx mori*, from days 0 to 7 of the last-instar larvae. For sample preparation, silk glands were extracted from one female and one male larva every 24 hours accurately after the fourth ecdysis. The reliability of these transcriptome data was confirmed by comparing the transcripts per million (TPM) values of the silk gene and quantitative reverse transcription PCR results. Hierarchical cluster analysis results supported the reliability of transcriptome data. These data are likely to contribute to the progress in molecular biology and genetic research using *B. mori*, such as elucidating the mechanism underlying the massive production of silk proteins, conducting entomological research using a meta-analysis as a model for lepidopteran insect species, and exploring medical research using *B. mori* as a model for disease species by utilising transcriptome data.

## Introduction

The domestic silkworm, *Bombyx mori*, is a lepidopteran insect renowned for silk production. Additionally, *B. mori* serves as a bioreactor, enabling the development of a baculovirus vector system with silkworm^1^ and a transgenic silkworm system^2,3^, leading to the production of recombinant proteins. Understanding the mechanisms regulating silk gene expression through its high-quality genome sequences is essential for improving the production of recombinant proteins. Therefore, *B. mori* draft genome sequence data were first published in 2004^4,5^ and have since been continuously updated with additional data^6,7^. Consequently, several related datasets, such as full-length cDNA data^8^, also have been published. These data are now available in public databases such as DDBJ/NCBI/ENA and in silkworm databases such as KAIKOBase, SilkBase, and SilkDB^9–11^. The development of these genome data and genetic tools has elevated the status of *B. mori* as a model lepidopteran species, contributing to advances in entomology (not limited to *B. mori*) and medical science^12,13^. To further expand the silkworm genome and related data, we constructed a chromosome-level reference genome and transcriptome datasets^7,14^. Using a new gene annotation workflow, we performed functional annotations on the silkworm gene^15^. Furthermore, we generated transcriptome expression data for multiple tissues in day-3 last-instar larvae (p50T (Daizo) strain, used for the genome project), when the silk gene was abundantly expressed^14^. The development of genome editing techniques for *B. mori* has enabled us to perform precise gene or genome modifications^16^. The *w1 pnd* strain, an experimental strain (non-diapause and white-eye) often employed for these modifications, is crucial for *B. mori* research.

The silk gland (SG) is a key tissue for silk production and cocoon formation, comprising three parts: the anterior silk gland (ASG), middle silk gland (MSG), and posterior silk gland (PSG); the MSG is further divided into the anterior part (MSG_A), middle part (MSG_M), and posterior part (MSG_P)^17,18^ (Fig. 1). Each SG component plays an important role in silk production and possesses a unique silk gene expression profile. During the life cycle of *B. mori*, the last-instar larvae produce silk by expressing silk genes, including *Sericin1, Sericin2, Sericin3, Fibroin-H, Fibroin-L, and fibrohexamerin (fhx)*. While *Sericin1, Sericin2*, and *Sericin3* are primarily expressed in MSG^18^, *Fibroin-H, Fibroin-L*, and *fhx* genes are expressed in PSG^19–21^. Considering these features, the time-course reference transcriptome expression data of MSG_A, MSG_M, MSG_P, and PSG from the last-instar larvae of the *w1 pnd* strain hold potentially high value and thus can serve as important reference datasets. Therefore, we conducted to construct these data. Actually, we prepared total RNA samples extracted from multiple SG territories from days 0 to 7 of the last-instar larvae (Fig. 1). After performing RNA sequencing (RNA-seq) on these samples, we calculated the expression values of the reference transcripts for each sample using pre-prepared RNA-seq data (trimmed and quality-controlled) and previously reported reference transcript data^14^. The expression data for each sample were merged into one matrix as the time-course expression data for SG (Fig. 2). These expression data are invaluable for various aspects of *B. mori* research, including the molecular mechanism of silk gene expression control, lepidopteran or insect basic research (e.g., comparative genomics), basic medical research (e.g., disease models using *B. mori*), and producing and breeding more valuable *B. mori* strains with enhanced silk production capabilities.

**Fig. 1.**
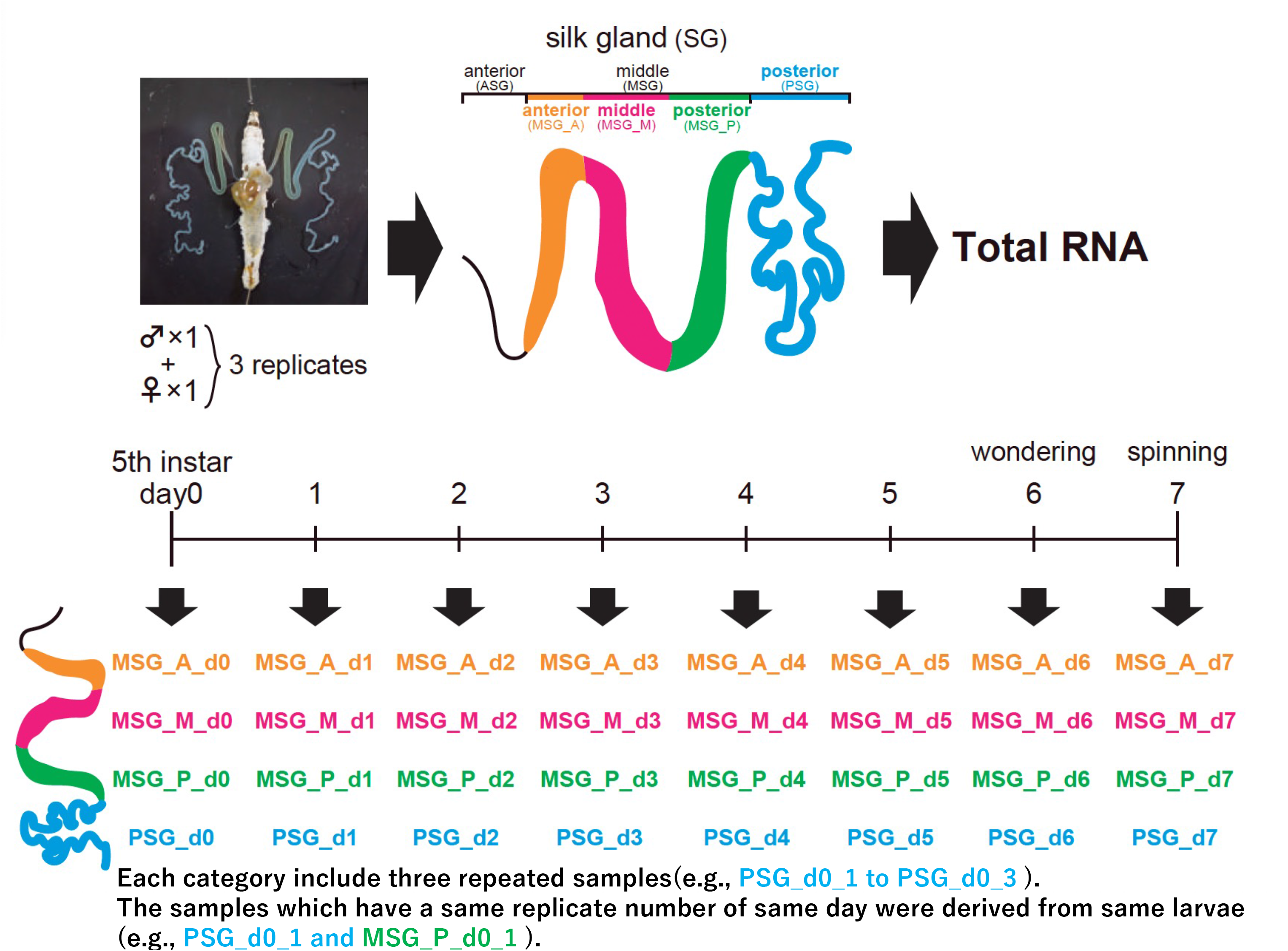
Schematic workflow of sample preparation for extraction of total RNA. Fifth-instar larvae, one female and one male, from day 0 to day 7, were used as one biological replicate with three replicates prepared. SGs were dissected from both larvae and separated into MSG_A, MSG_M, MSG_P and PSG. Female and male dissections were combined to form one sample. Total RNA was extracted from each prepared sample (upper panel). Abbreviations of sample names in this work are shown in the lower panel.

**Fig. 2.**
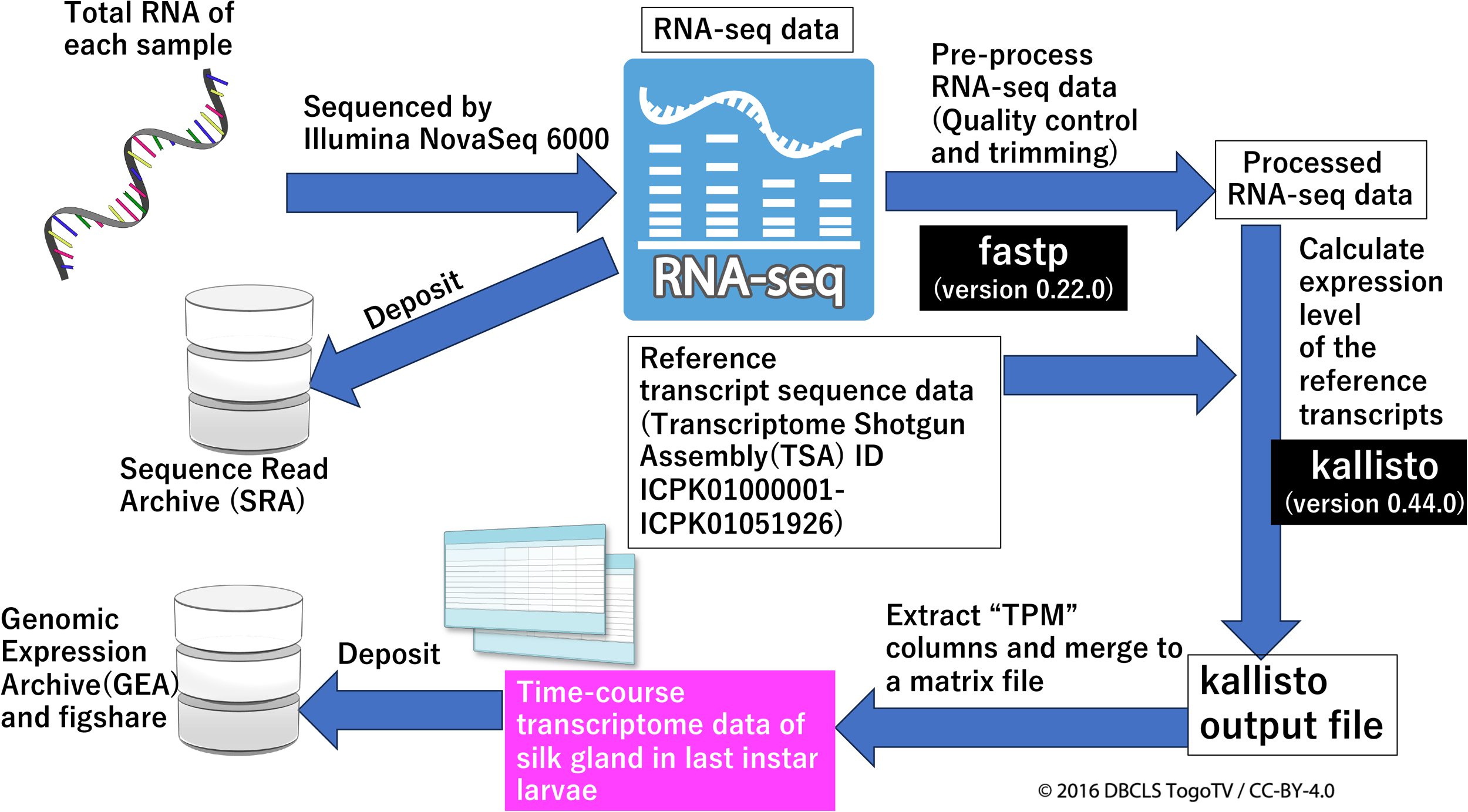
Schematic workflow of transcriptome analysis. Total RNA from the samples shown in Figure 1 was sequenced. The raw RNA-seq data were deposited into SRA. Quality control and trimming of the raw RNA-seq data were performed using fastp. Calculations of the reference transcripts of each sample were performed using processed RNA-seq and reference transcript data by kallisto. The expression data from each sample (Supplementary File 2) were merged into a matrix data (“Time-course transcriptome data of the silk gland in the last-instar larvae”). These expression data were deposited in the GEA and figshare (Supplementary File 3).

## Materials and Methods

### Sample preparation and total RNA extraction

The *w1 pnd* strain of *B. mori* silkworms was raised on an artificial diet (Nihon Nosan Kogyo, Yokohama, Japan) at 25 °C under 12:12 h light/dark conditions. Immediately after the fourth molting, the larvae were individually placed in plastic petri dishes containing the artificial diet. Every time a larva moulted during the last-instar period, one male and one female insect were prepared and maintained for 24 h (days 0–7). Subsequently, SGs were dissected individually from the prepared male and female larvae in a physiological saline solution. The extracted SGs were further dissected into MSG_A, MSG_M, MSG_P, and PSG. Each SG part was then homogenised in a TRIzol (Invitrogen, Carlsbad, CA, USA) solution and stored at -80 °C until all sample series were collected. Chloroform was added to the above solution, and the supernatant after vortexing and centrifugation was mixed in equal volumes (300*μ*L) from both sexes to make the primary solution. Total RNA was extracted from primary solution with the removal of genomic DNA by the RNeasy Plus Mini Kit (Qiagen, Hilden, Germany). The purity and concentration of extrasted totel RNA were confirmed using a NanoDrop ONE spectrophotometer (Thermo Fisher Scientific, Waltham, MA, USA). Three-biological replicates were prepared for each sample category (e.g., MSG_A_d0_1 to MSG_A_d0_3) using the different larvae (Fig. 1).

### Library preparation and RNA-seq

For RNA-seq, cDNA libraries were constructed from the total RNA samples using the TruSeq Stranded mRNA Library Prep Kit, following the procedures outlined in its Reference Guide. Sequencing was performed using an Illumina NovaSeq6000 (San Diego, California, USA.). Both library construction and sequencing were performed by Macrogen Japan Corp (Kyoto, Japan).

### Data analysis for transcriptome data

The workflow for analysing the transcriptome data is shown in Fig. 2. All raw RNA-seq sequence data (FASTQ files) have been deposited in the Sequence Read Archive (SRA) (for details, see Supplementary File 1 and Data Records) (Fig. 2). Pre-processing of the raw sequence data involved removing adapter sequences, trimming, and quality control using fastp (version 0.22.0) with default settings^22^. Subsequently, using processed RNA-seq data and reference transcript sequence data of *B. mori* (Transcriptome Shotgun Assembly (TSA) accession ID ICPK01000001-ICPK01051926)^14^, TPM values of the reference transcripts in each sample were calculated using kallisto (version 0.44.0) with a bootstrap value of 100 and others as defaults^23^. A TPM column in abundance.tsv of each sample, which is a kallisto output data file (Supplementary File 2), was extracted, which contains time-course transcript data of each sample. Another column data including transcript IDs of the reference transcript were extracted from abundance.tsv of one sample. Finally, the time-course data for all samples and the transcript ID data were merged into a single matrix dataset as “Time-course transcriptome data of the silk gland in *B. mori* last-instar larvae” (Supplementary File 3). The matrix data were deposited in the Genomic Expression Archive (GEA) of the DNA Data Bank of Japan (DDBJ) (Accession ID: E-GEAD-662, URL: https://www.ddbj.nig.ac.jp/gea/index-e.html) (Fig. 2). All the commands used in these analyses are available in Supplementary File 4.

### Hierarchical clustering analysis

To assess the similarities in expression profiles among all samples, hierarchical clustering analysis was performed using R (version 4.2.3) in RStudio (version 2023.03.0+386) with expression matrix data as input. Spearman’s rank correlation coefficient and Ward’s method were used for measuring similarities between each sample and clustering, respectively. The R code for this analysis is uploaded in Supplementary File 4.

### Data Records

The RNA-seq raw data in this study were deposited in the SRA under BioProject ID PRJDB16887. Accession IDs of BioSamples and SRA for each RNA-seq sample are described in Supplementary File 1. Additionally, expression data across the reference transcript in all samples can be accessed via GEA (Accession ID: E-GEAD-662) or figshare (Supplementary File 3).

### Code Availability

Code files for transcriptome analysis and hierarchical clustering analysis using R, as shown in Fig. 2, are available in Supplementary File 4.

## Results and Discussion

Using RNA-seq data and reference transcriptome data^14^, we calculated TPM values of each transcript in ASG, MSG_A, MSG_M MSG_P and PSG of last instar larvae from days 0-7 (Fig. 2). The calculated TPM values were in one matrix data (Supplementary File 3). To assessed reliability of the constructed time-course transcriptome data, we performed comparison of representative silk genes of TPM values with qPCR results and hierarchical clustering analysis.

### Transcriptome data validations using TPM values of silk genes compared to the silk genes expression results by qPCR

To confirm the reliability of the expression values, we compared the time-course TPM values of representative silk genes with the relative expression values using quantitative reverse transcription PCR (RT-qPCR) from the total RNAs^24^. The TPM expression values of *Sericin1, Sericin2, Sericin3, Fibroin-H, Fibroin-L*, and *fhx* are shown in Fig. 3a–f, respectively. In Fig. 3a, the TPM time-course values (total TPM values of all isoforms) of *Sericin1* (Transcript ID: KWMTBOMO06216, MSTRG.2477.1-MSTRG.2477.16) in MSG_A are shown in the upper left graph. Although TPM values exhibited variations in larvae expressing *Sericin1*, this gene began to be expressed in day-2 larvae, peaked in day-3 larvae, and decreased after day 4. *Sericin1* expression in MSG_M increased and peaked on days 4 and 5 and decreased in day 6 larvae (lower left graph in Fig. 3a). These features were consistent with the qPCR results reported previously^24^, except for the timing of the decline, as *Sericin1* expression decreased from day 5 larvae onward in the qPCR results. The expression profile of TPM values of *Sericin1* showed an increase in MSG_P from days 0 to 5 and plateaued afterward (upper right graph in Fig. 3a), whereas *Sericin1* qPCR results showed that the mRNA levels of *Sericin1* in MSG_P increased from day 0 to day 2 larvae and plateaued afterward. Interestingly, the time points at which *Sericin1* expression levels plateaued in the MSG_P group differed between the TPM and qPCR results^24^. These differences in the TPM and qPCR profiles may be because the qPCR values cannot contain all the isoforms described above with the primer pair used. Notably, *Sericin1* TPM values in MSG_P and MSG_M from day 0 to day 7 larvae were notably higher than those in MSG_A, as observed in the qPCR results (lower right graph in Fig. 3a). *Sericin1* TPM values in MSG_P from day 5 to day 7 were larger than those in MSG_M. However, this pattern was not observed in the qPCR results. The relative mRNA values in MSG_M were larger from day 0 to day 7 (but not significantly different)^24^. These discrepancies could be due to the methods used to measure gene expression values. As described above, TPM values include all isoforms, whereas qPCR values may not. *Sericin1* mRNA levels in MSGs from day 0 to 7 larvae (*w1 pnd* strain) using qPCR showed that the amounts of *Sericin1* mRNA increased from day 0 to day 6 and decreased on day 7, which was comparable to the MSG_P profile of the TPM results of this work^2^. The TPM values of *Sericin2* (Transcript ID: KWMTBOMO06334, MSTRG.2627.1, and MSTRG.2627.2) in MSG_A of larvae from day 0 to day 4 were notably high (over 15000) and dropped to nearly zero from day 5 to day 7 (upper left graph in Fig. 3b). In MSG_M, *Sericin2* TPM values decreased from approximately 11000 on day 0 to low levels in day 3 larvae and remained very low in day 7 larvae (lower left graph in Fig. 3b). Similarly, TPM values of *Sericin2* in MSG_P were over 11000 in day 0 larvae, dropped below 1000 in day 1 larvae, and approached nearly zero in larvae of days 2 to 7 (upper right graph in Fig. 3b). Comparing the TPM values across samples tested, *Sericin2* TPM values were the highest in day 0 larvae (MSG_A having the highest value and MSG_P the lowest), decreasing dramatically across larval stages (lower right graph in Fig. 3b). These trends of *Sericin2* TPM values were consistent with the qPCR results ^24^ and in a previous report^18^. *Sericin3* (Transcript ID: KWMTBOMO06311 and MSTRG.2595.1-MSTRG.2595.9) TPM values in MSG_A were negligible in day 0 larvae and gradually increased in day 2 to day 4 larvae (ranging between 3000 and 65000). Subsequently, the TPM values dramatically increased in day 4 and day 5 larvae, plateauing in day 5 to day 7 larvae (upper left graph in Fig. 3c). In MSG_M, *Sericin3* TPM values increased in day 1 to day 5 larvae, with a sharp increase in day 4 to day 5 larvae, followed by plateauing until day 7. The MSG_P, *Sericin3* TPM values were very small or nearly zero at all stages tested (lower left and upper right graphs in Fig. 3c). *Sericin3* TPM values in MSG_A and MSG_M from days 3 to 7 were notably higher than those in MSG_P (lower right graph in Fig. 3c), consistent with qPCR results^24^. In previous studies, *Sericin3* expression profiles revealed that *Sericin3* expression was only observed in MSG_A of days 5 to 7 larvae, with significantly higher expression observed in the day 6 and day 7 larvae compared with that in day 5 larvae^18^. These findings are consistent with the results of TPM data from our study. Specifically, *Sericin3* TPM values in MSG_A of day 5 to day 7 larvae were notably higher than those in the other samples. In PSG, *Fibroin-H* (Transcript ID: KWMTBOMO08464 and MSTRG.14927.1-MSTRG.14927.23) TPM values increased from day 0 to day 2, remained relatively stable from day 2 to day 5, and slightly increased from day 5 to day 7 (Fig. 3**d**), whereas *Fibroin-L* (Transcript ID: KWMTBOMO15365 and MSTRG.5511.1) TPM values gradually increased from day 0 to day 6 and plateaued (Fig. 3e). The expression profiles of *Fibroin-H* and *Fibroin-L* TPM values from transcriptome analysis were similar to those determined by qPCR^24^. *Fhx* (Transcript ID: KWMTBOMO01001 and MSTRG.10154.1) TPM values increased from day 0 to day 5 and decreased and plateaued from day 5 to 6 and from day 6 to 7, respectively (Fig. 3f). Fhx is associated with Fibroin-H and Fibroin-H linked by disulfide bond^19^, suggesting that the *Fhx* expression profile was similar to the profiles of *Fibroin-H* and *Fibroin-H*. The expression profiles of these three genes determined based on the TPM values were similar. Taken together, these comparisons of the six silk gene expression data using TPM values in this study, qPCR results, and findings from other reports suggest that the TPM transcriptome expression data in this study were reliable.

**Fig. 3.**
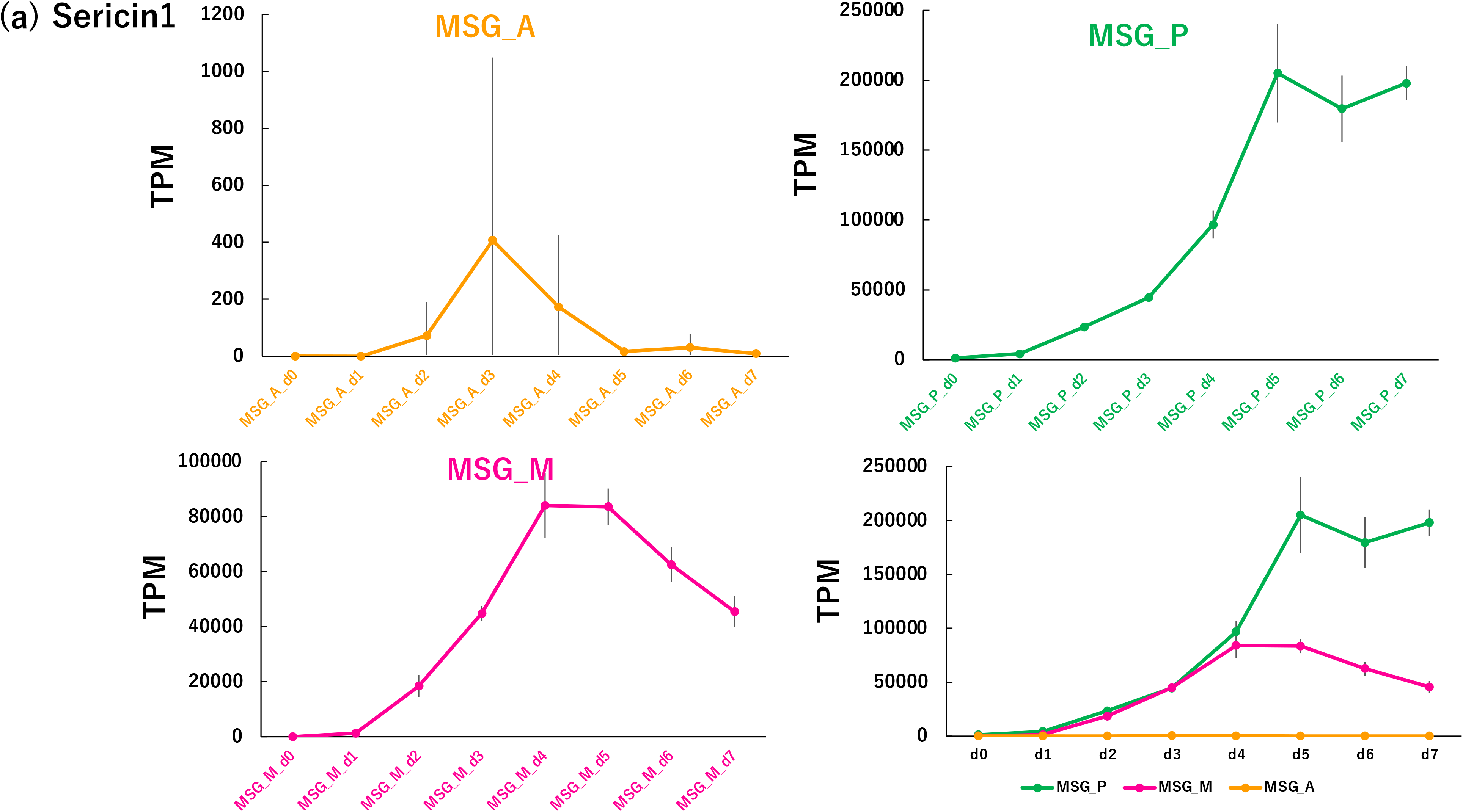
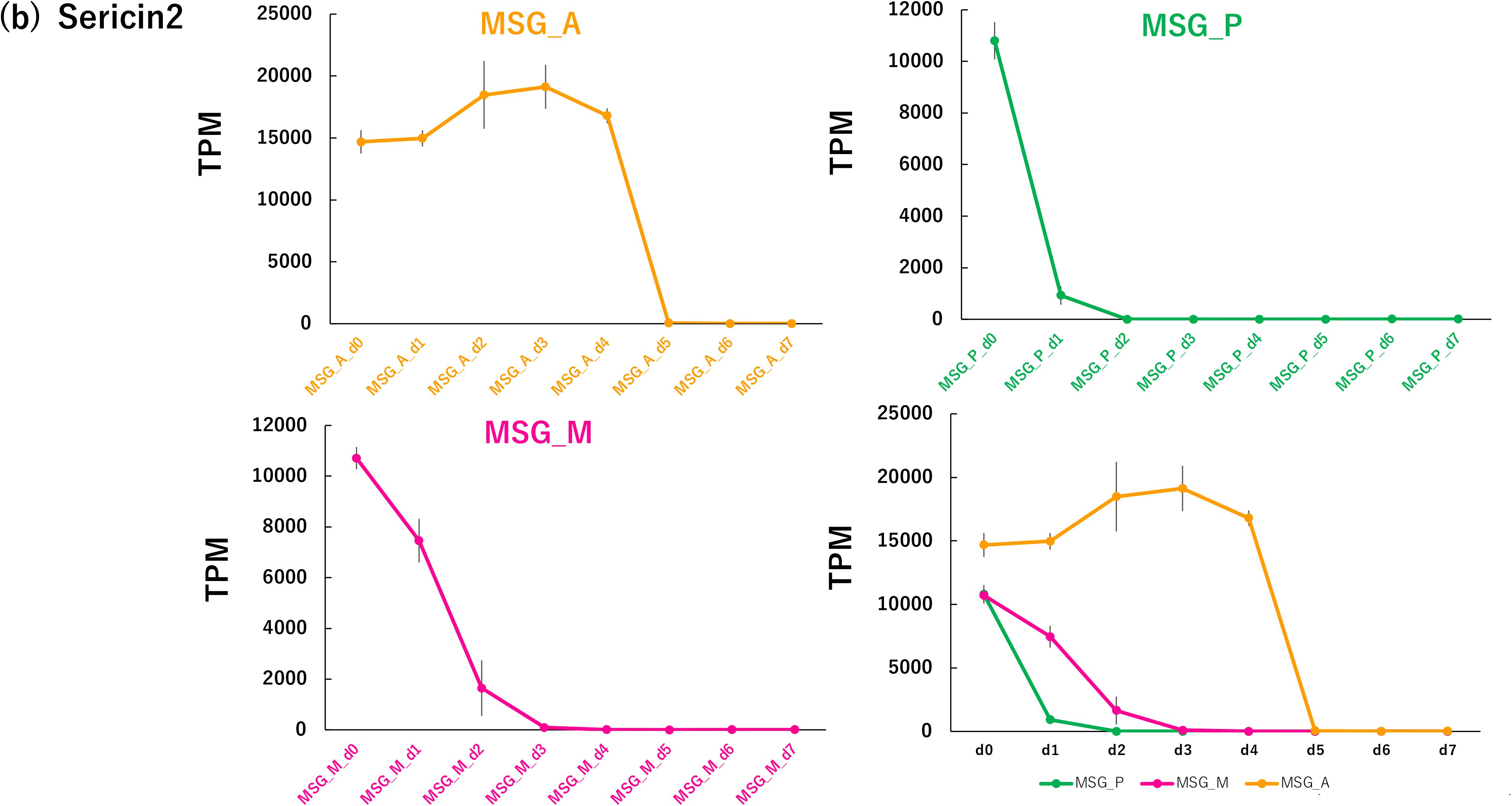
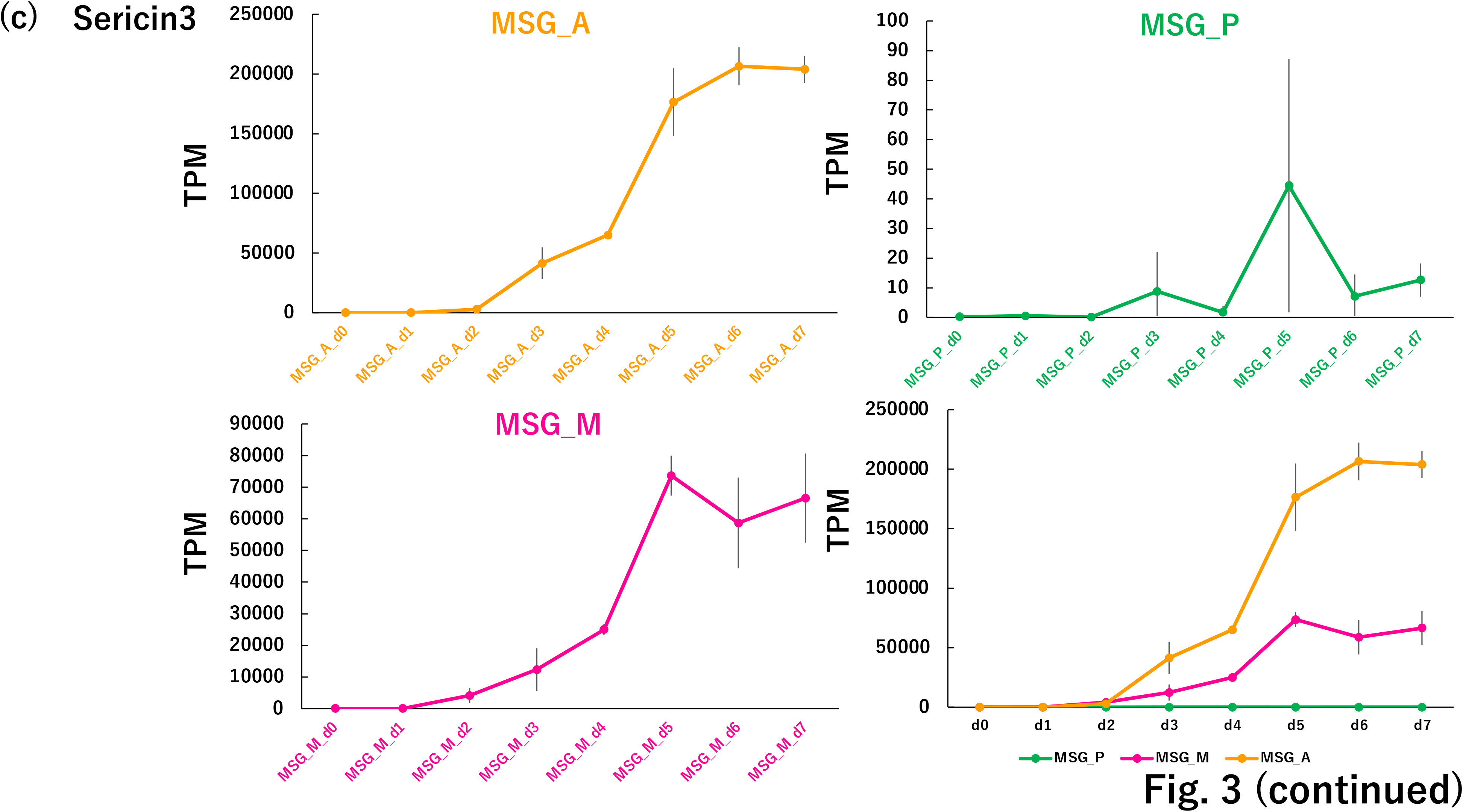
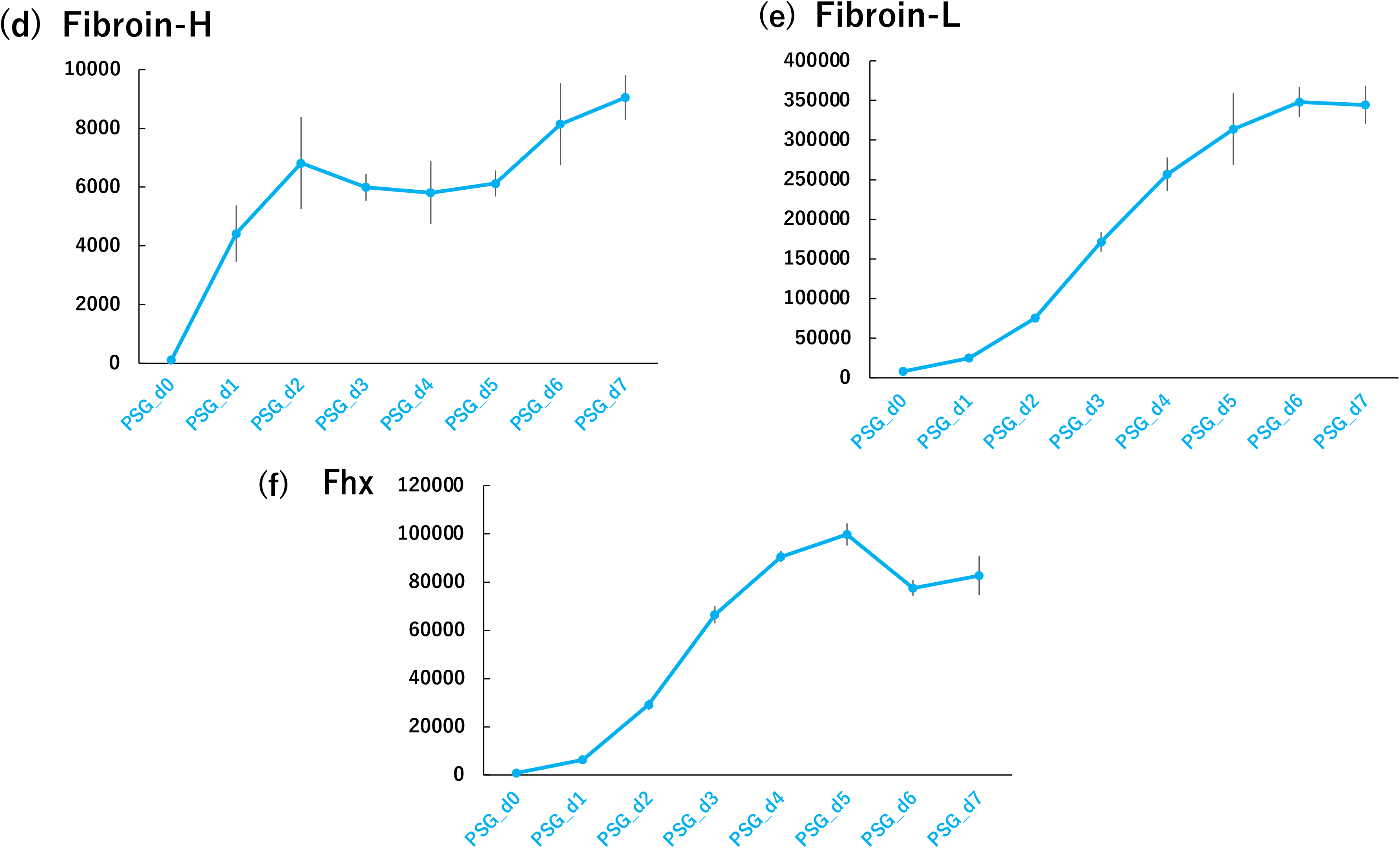
Expression profiles of the major silk genes. Average TPM values representing the total TPM values of all isoforms for six silk genes per biological replicate were calculated (See main text). *Sericin1* (a), *Sericin2* (b) and *Sericin3* (c) expression levels in MSG_A (left upper graph), MSG_M (left lower graph), MSG_P (right upper graph) across day 0 to day 7 last-instar larvae (three replicates) are presented. The right lower graph displays the average TPM values of the three SG samples. Average TPM values of *Fibroin-H* (d), *Fibroin-L* (e) and *Fhx* (f) in PSG are shown. Total TPM values of all isoforms for each silk gene per biological replicate were calculated. For example, total TPM value of *Sericin1* of each biological replicate was calculated by using all TPM values of *Sericin1* isoforms within that biological replicate. The vertical bar represents the standard deviation of each point.

### Transcriptome data validations by hierarchical clustering analysis

Transcriptome data validation was performed through hierarchical clustering analysis using the TPM matrix data^25^ as input. The results are depicted in Fig. 4. The dendrogram results show that the three biological replicates of all samples, except one (MSG_P_d0 samples), form a single cluster or small closely neighbouring clusters. Specifically, MSG_P_d0_3 is located in the MSG_M_d0 cluster. We presumed that this was because the MSG_P_d0_3 sample contained a small number of MSG_M_d0 cells, and the contaminated MSG_M_d0 cells, which expressed a large number of genes, masked the transcriptome features of MSG_P_d0. If this is true, the samples are placed in different clusters. Typically, most samples from the same category, such as MSG_M, MSG_P, and PSG, clustered together or formed two distinct intermediate clusters (MSG_A). MSG_A_d0-4 and MSG_A_d5-7 samples were located in two separate intermediate clusters. The differences in the certain gene expressions, particularly evident in highly expressed genes such as *Sericin*2 and *Sericin3* (Fig. 3), may reflect the two separate intermediate clusters. Several samples, including PSG_d0, MSG_P_d0-2, and MSG_M_d0-2, grouped together in clusters, possibly reflected similarities in the abundantly expressed genes with similar expression profiles across these samples rather than the same territories of SG. Indeed, *Sericin1* and *Sericin3* TPM values between MSG_P_d0-2 and MSG_M_d0-2 were not significantly different from those of the other samples (Fig. 3). Additionally, *Fibroin-H, Fibroin-L* and *Fhx* expression levels in PSG_d0 were markedly lower than those in the other PSG samples. In summary, the hierarchical clustering analysis demonstrated that, except for a few samples, there were no significant differences in transcriptome TPM values between biological replicates, suggesting that the reliability of the transcriptome data in this study.

**Fig. 4.**
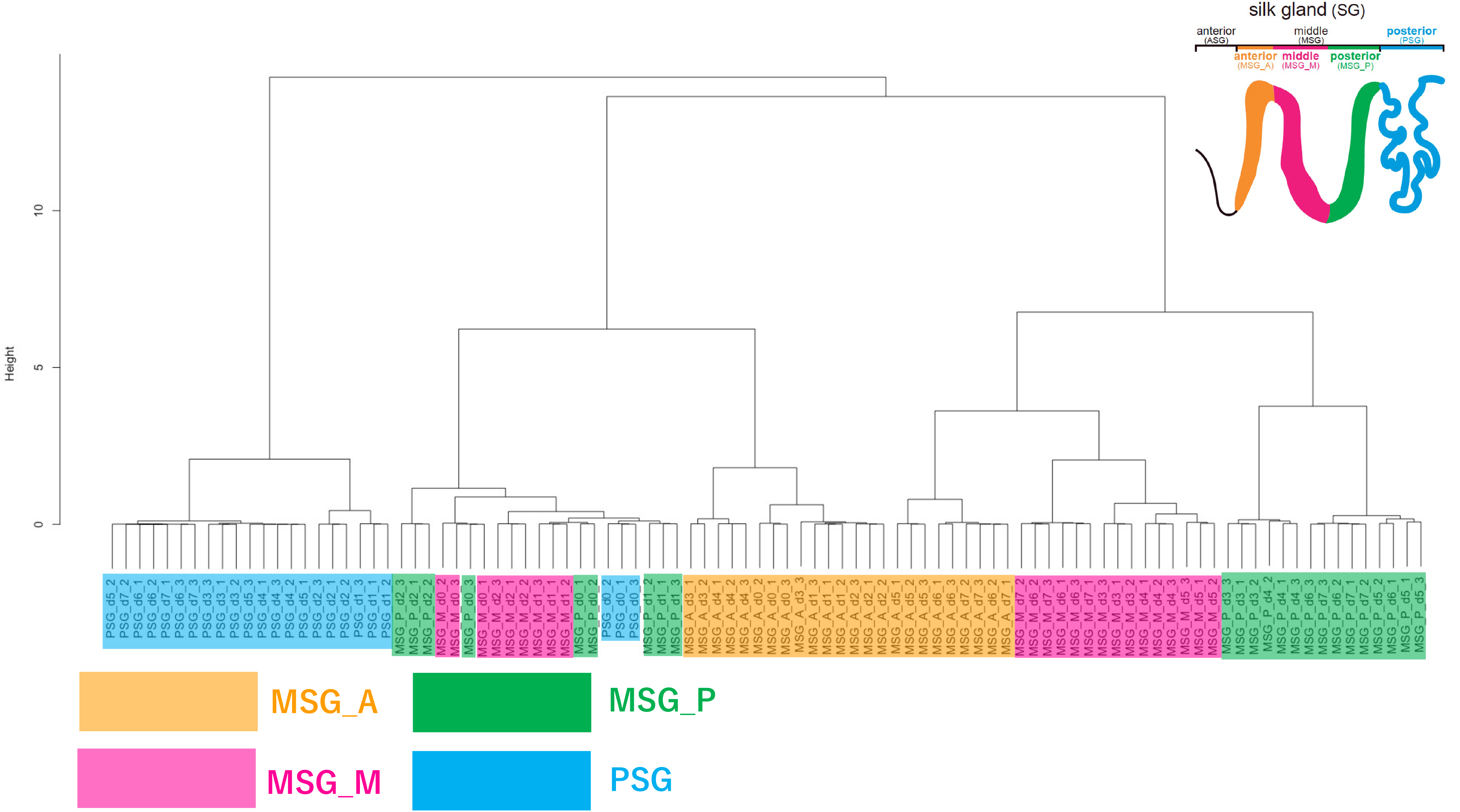
Dendrogram of hierarchical clustering analysis. Hierarchical clustering analysis using time-course transcriptome data (Supplementary File 3) as input was performed. The resulting dendrogram was constructed based on the results of hierarchical clustering analysis. MSG_A, MSG_M, MSG_P and PSG samples in the dendrogram are coloured yellow, purple, green, and blue, respectively. A schematic representation of the SG (as shown in Fig. 1) is included in the upper right corner.

In conclusion, two data analysis described above showed that time-course transcriptome data of silk glands possess the reliability, which can be utilized for the multiple studies.

## Acknowledgements

This work was supported by the MAFF Commissioned project study on the “Research project for sericultural bio-industry” (Grant Number JP22680575) to Y.M., T.T., H.S. and K.Y., ROIS-DS-JOINT (024RP2023) to Y.M, A.J., H.O., H.C., H.B. and K.Y., and the Center of Innovation for Bio-Digital Transformation (BioDX), an open innovation platform for industry-academia co-creation (COI-NEXT) of JST (COI-NEXT, JPMJPF2010) to Y.M., A.J., T.T, H.S., H.B. and K.Y. Parts of the Fig. 2 were drawn by using illustrations from TogoTV (© 2016 DBCLS TogoTV, CC-BY-4.0 https://creativecommons.org/licenses/by/4.0/deed.ja).

## Author contributions

Y.M., H.O., H.C., H.S., H.B. and K.Y. conceived the study. Y.M. prepared the RNA samples from silkworms for RNA-seq. Y.M., A.J., H.B., and K.Y. performed the bioinformatics data analysis and data registration. All authors curated and validated the data and metadata analysed in this study. K.Y wrote the original draft of the manuscript. All authors reviewed and edited the draft of the manuscript. H.O., H.S., H.B. and K.Y. contributed to funding acquisition. All the authors have read and agreed to the published version of the manuscript.

## Supplementary Information

All supplementary files are available in figshare (DOI: 10.6084/m9.figshare.c.6978654).

**Supplementary File 1**

List of metadata for all RNA-Seq data. The sample description, Biosample ID, SRA Accession ID (SRA run ID), and Fastq file names of each RNA-seq dataset used in this study are listed (DOI: 10.6084/m9.figshare.24921426).

**Supplementary File 2**

Kallisto output files. PSG_kallisto.tar.gz, MSG_P_kallisto.tar.gz, MSG_M_kallisto.tar.gz, and MSG_A_kallisto.tar.gz are collections of kallisto output files of the prepared samples. samle_name_directory_name_list.xlsx is the list of samples and Kallisto output directory names (DOI: 10.6084/m9.figshare.24978483).

**Supplementary File 3**

Matrix data of the expression values of all reference transcripts in all samples. Expression values (TPM values) of reference transcripts were calculated using all RNA-seq data, and all expression values are summarized in the matrix data (DOI: 10.6084/m9.figshare.24921435). These data are available in the DDBJ GEA (Accession ID: E-GEAD-662).

**Supplementary File 4**

Script file for transcriptome and hierarchical clustering analyses. The fp_ks_script.sh file contains codes of fastp for pre-processing the raw sequence data and kallisto to calculate the expression values of each sample. The TPM_script.sh file contains commands for constructing transcriptome TPM matrix files from the kallisto output files (Supplementary File 2). “script_r.sh” is a file for performing hierarchical clustering analysis (DOI: 10.6084/m9.figshare.24921459).

